# BLIMPS: a technique for tandem biosensor imaging across multiple populations of presynaptic terminals, using lattice light sheet microscopy

**DOI:** 10.64898/2026.02.12.705649

**Authors:** Mariana Potcoava, Zack Zurawski, Iris Lu, Simon Alford

**Affiliations:** Department of Anatomy and Cell Biology, University of Illinois at Chicago, 808 South Wood Street, Chicago, IL 60612, USA

## Abstract

Within neuronal circuits, ordered neurotransmission is contingent upon balance between excitatory glutamatergic and inhibitory GABAergic signaling. To study circuit-level processes, the paradigm of 4D cellular physiology has been developed, where, single cells and subcellular structures are studied as individual units in three-dimensional space over a continuous interval rather than as a single moment in time, or as a population-level average. Neurons are excitable cells expressing voltage-gated Ca^2+^ channels and Ca^2+^ fluxes subsequent to action potential firing are widely used as markers of neuronal activity. While the imaging of Ca^2+^ dynamics at the soma is often performed, the imaging of Ca^2+^ fluxes at presynaptic terminals has often proven to be an experimental challenge: existing imaging modalities suffer from inadequate acquisition speeds, insufficient penetration depths, insufficient spatial resolution to identify axonal structures, or spectral crosstalk issues. To visualize presynaptic Ca^2+^ dynamics in both excitatory and inhibitory neurons, here we combine advanced lattice light-sheet microscopy with viral delivery of two genetically encoded calcium indicators (GECIs)- jRGECO1a and jGCaMP8f, to perform sequential imaging of Ca^2+^ dynamics within acute *ex vivo* slice preparations. Our methodology, Biosensor Lattice light-sheet Imaging of Multidimensional Presynaptic Structure (BLIMPS), includes acute brain slice preparation, mounting on a temperature-controlled flow chamber within a LLSM, and imaging of electrically evoked Ca^2+^ signals, with high adaptability to a range of genetic and pharmacological disease models. Our technique offers high spectral separation between evoked signals from each of the two GECIs and fast acquisition speeds of 0.1-0.3 KHz. Included within the BLIMPS technique is a robust, open-source data analysis pipeline to track highly responsive neuronal structures such as presynaptic terminals and quantify both the amplitudes and decay rates of evoked fluxes.

## INTRODUCTION

Many physiological processes are contingent on antagonistic relationships between discrete cellular populations, forming precise control from negative feedback loops. In the brain, neuronal homeostasis is dependent on a balance between excitatory and inhibitory (E-I) neurotransmission, and E/I imbalance has been linked to progressive conditions such as Alzheimer’s disease^1,2^: Early imbalances towards excitation precede later loss of synapses and soma^3^. The E-I ratio has traditionally been assessed as a ratio of excitatory and inhibitory postsynaptic currents using electrophysiology^4-7^: this method provides well-validated readouts with very high temporal fidelity (tens of KHz) from whole neurons, but has minimal capability to differentiate between defects in axonal, dendritic, and somatic structures, the whole cell patching technique itself may interfere with neuronal physiology, and assays of presynaptic activity are only available as indirect functions of postsynaptic ligand-gated ion channels such as AMPARs.

Recent advances in fluorescence microscopy and genetically encoded biosensors empower researchers with the tools to directly image cellular processes such as presynaptic Ca^2+^ fluxes with meaningful spatiotemporal resolution. Imaging allows for the visualization of select physiological structures and processes without extensive disruption of cellular function and can provide higher throughput readouts: many cells can be tracked independently in one field of view, while patch-clamp electrophysiology is limited to a small number of cells.

Optical physics has pushed spatiotemporal resolutions past the diffraction limit of light and to kHz frame rates, attempting to capture cellular processes as they happen in real-time. Scanning microscopy techniques such as STED can image neuronal structures smaller than the diffraction limit of visible light^8^. While super-resolution microscopy has been developed to image at resolutions down to 10nm^9^, this technique is most useful for imaging tissue at a depth of 20-50μm, along with slow acquisition times, photobleaching, and phototoxicity from the high-intensity laser illumination^9,10^, limiting its utility to capture processes like neurotransmission that happen at micro- to millisecond speeds, relegating these techniques to structural or fixed tissue imaging^10^. The development and widespread implementation of multiphoton imaging in the 21^st^ century has revolutionized the imaging of genetically encoded biosensors, offering penetration depth of up to 1mm at acquisition speeds of approximately 30 Hz^11^, with specialized implementations offering faster kinetics for smaller fields of view^12^. These systems are widely used for in vivo imaging of brain tissue in live animals^13,14^, permitting researchers to correlate biosensor dynamics with behavior. However, spectral crosstalk is a major barrier to simultaneous imaging of multiple fluorophores in multiphoton microscopy: multiphoton excitation spectra are broad, complex, and often exhibit multiple peaks^15^, while traditional single-photon imaging systems can use well-defined laser lines to excite each fluorophore with minimal crosstalk. Light-sheet microscopy technologies utilize a thin sheet of laser light to illuminate only a narrow(<1μm) plane of a sample, minimizing photobleaching while retaining high spatial resolution^16^. The advent of lattice light sheet microscopy (LLSM) systems by Eric Betzig in 2014 offers unparalleled potential for fast biosensor imaging^17^: in these systems, a spatial light modulator transforms coherent light into shaped, non-diffractive Bessel beams, eliminating the high-energy outer rings through destructive interference. Uniform illumination is maintained throughout the x-axis by rapid oscillation from a galvanometer. These methodologies can achieve transverse resolutions of approximately 230 nm and axial resolutions of 370 nm^17^.

Because the entire plane is imaged simultaneously, these systems can reach acquisition speeds up to 0.30 kHz while still resolving distinct synaptic puncta and cellular structures^18-20^, making it ideal for imaging of neuronal Ca^2+^ dynamics in live tissues. The time course of neuronal Ca^2^+ fluxes has been estimated to be in the single-digit millisecond range via the use of organic Ca^2+^-sensitive dyes^21,22^, indicating that LLSM has adequate temporal resolution to visualize this process.

The introduction and continuous improvement of genetically encoded calcium indicators (GECIs) has been of massive utility to the field of live cell imaging. Originally developed as a fusion of calmodulin, circularly permuted green fluorescent protein, and the M13 fragment of myosin light chain kinase, the GCaMP family of probes becomes more fluorescent in response to Ca^2+^ binding^23^. After two decades of protein engineering, current variants such as jGCaMP8f offer ultra-fast kinetics, with rise times of 2ms and a 1.4-fold increase in fluorescence upon Ca^2+^ binding^24^. These sensors were accompanied by red GECIs such as jRCAMP1a and jrGECO1a^25^, which were originally intended to take advantage of the deeper penetrating capabilities of longer wavelengths of light in *in vivo* imaging. The non-overlapping excitation and emission spectra of jGCaMP and jRGECO1a allow simultaneous two-channel imaging without crosstalk, enabling independent monitoring of distinct calcium pools. The expression of genetically encoded sensors can be directed to specific cells with heterologous expression systems that feature cell-type specific promoters such as AAVs. By targeting each indicator to specific subcellular domains, such as cytosol versus mitochondria, or to different neuronal populations, researchers can map parallel signaling pathways in real time. Here, we introduce BLIMPS, a dual-color strategy transforming calcium imaging into a multidimensional analysis.

## RESULTS

### Performance Metrics of LLS

For BLIMPS, we built an LLS instrument utilizing existing specifications from the laboratory of Eric Betzig (Janelia Research Campus)^17^ (Figure 1A). To fully characterize the performances of the LLS system, we analyzed the modulation transfer function (MTF) and point spread function (PSF) of fluorescent latex beads (500 nm, λ_exc_ = 488 nm, Molecular Probes, Waltham, MA, USA), (Figure 1B). We prepared a bead solution (2% solids) diluted with distilled water (1:4000). Thin layers of bead solution were applied on coverslips. After drying, the cover slip was mounted on the sample holder under distilled water. Volumetric imaging of beads was performed using the LLS, Figure 1a. The scanning area is localized in the middle of the camera FOV (red dashed rectangle in the upper left corner of Figure 1a), and the Bessel beams cannot excite beads outside that area. We then determined the performance metrics’ distribution, MTF (Figure 1C) and PSF(Figure 1D), of the #2 bead, localized at z-galvo 0 μm.

**Figure 1.**
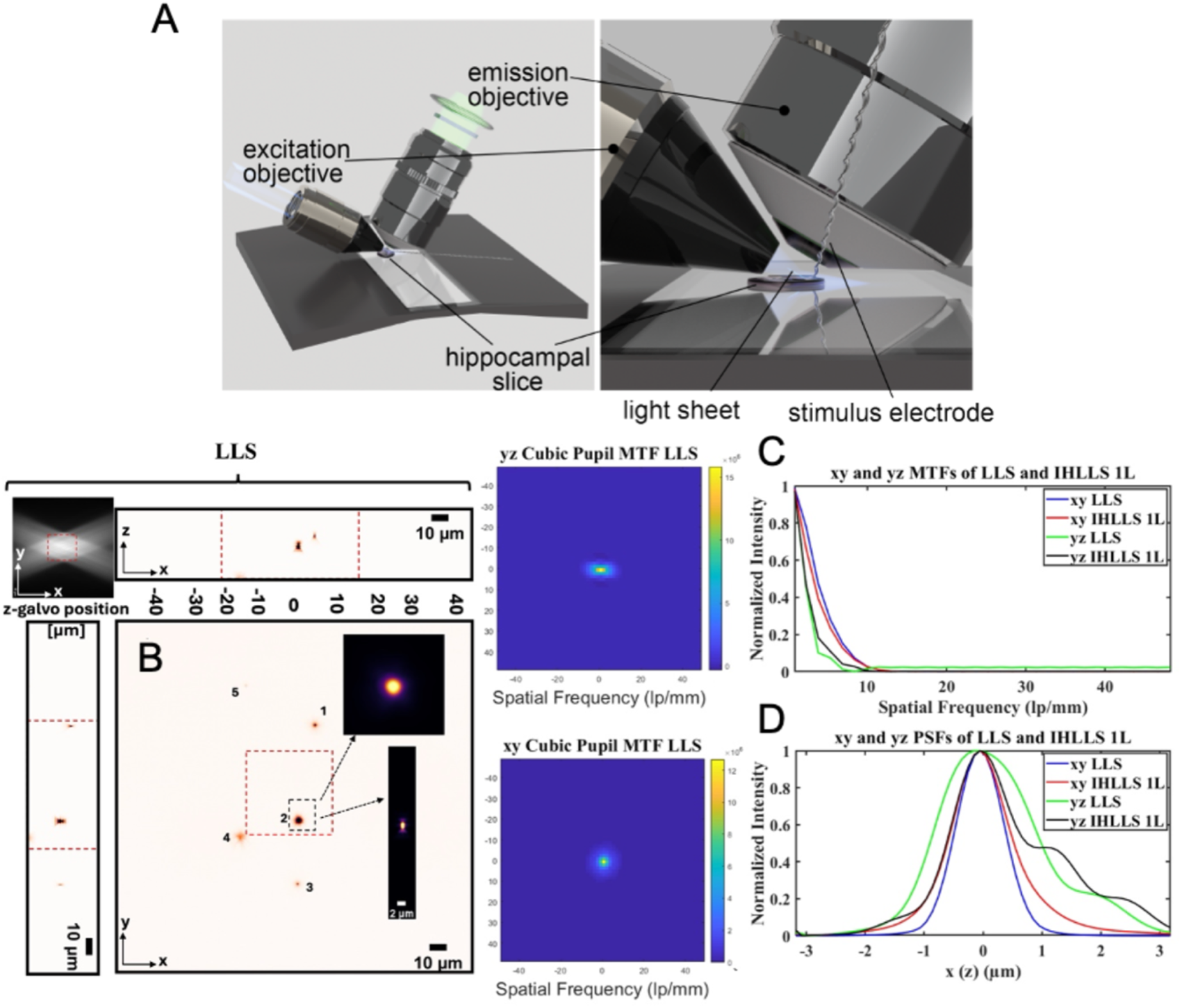
Axial and transverse resolution of the BLIMPS system. A. Diagram of acute hippocampal slice and custom PMMA stage in relation to lenses. B. Tomographic imaging of 0.5 µm fluorescent latex beads (500 nm, λ_exc_ = 488 nm), FOV 208 µm^2^, in the LLS system without deconvolution. On the sides and above are shown the max projections through the volume (400 Z-galvo steps). The Bessel beams are displayed in the upper left corner of each xy-projection to show the orientation of the beams (FOV 208 µm^2^). The red area enclosed inside the colored dashed rectangles is the scanning area for the LLS (52 µm^2^). The bead #2 in the black dashed rectangle that is in the middle of the lattice sheet is considered for calculating the resolution for the instrument. C. The axial MTFs of the LLS imaging techniques is shown in and the transverse MTFs.,. D. 1D xy and yz sections of the MTFs, 1D xy and yz of the PSFs. The FWHM of the curves are blue-0.5010 µm, green 0.8341 µm. The beads at λ_exc_ = 560 nm have similar behaviour.

### Spectral separation of jrGECO1a and jGCaMP8f fluxes

To determine if we could separately transduce excitatory glutamateric and inhibitory GABAergic neurons, we co-injected AAV5-mDlx-jGCaMP8f-WPRE and AAV(DJ)-CaMKII-NES-jRGECO1a-WPRE into the CA1 region of mouse hippocampi (Figure 2A). After a latency period of approximately six weeks for expression, acute hippocampal slices were prepared and the expression of jGCaMP8f and jRGECO1a was assessed, first utilizing a 40x epifluorescence microscope. Imaging of yellow 560nm fluorescence subsequent to excitation with a green LED (530 ± 10 nm) showed fluorescent axons throughout the stratum oriens and stratum radiatum(Figure 2Bi-Biii), while imaging of green 535nm fluorescence subsequent to illumination with a blue LED (470 ± 15 nm) showed a small number of fluorescent soma forming a dense mesh around the stratum pyramidale (Figure 2Bii-Biii). After butyl cyanoacrylate mounting to a glass coverslip, LLSM imaging of these regions with BLIMPS showed clear non-overlapping fluorescent axons, as well as accumulation of both GECIs, but particularly jRGECO1a, in puncta, consistent with prior reports of accumulation of fluorescent proteins in lysosomes^25^. Minimizing spectral crosstalk is a requirement for any multichannel imaging technique. To assess the extent of spectral crosstalk between 535nm and 590nm fluorescence, we assessed the pixel-by-pixel spatial correlation (Figure 2C) of each channel at 3 positions on the x-axis-the 64^th^, 256^th^, and 448^th^ scanlines, at lattice penetration depths of 5 and 18μm (Figure 2Ci-2Cii) from the surface of the slice. At a penetration depth of 5 μm, the 64^th^ scanline near the smallest x-coordinate, the 535nm and 595nm channels had a positive correlation of 0.623±.048, which diminished to 0.536±.045 and 0.365±0.071 for the 256^th^ and 448^th^ scanlines. In contrast, at the deeper penetration depths of 18 μm, the 64^th^ scanline had a correlation of 0.320±0.077, the 256^th^ scanline had a correlation of 0.344±0.0762, and the correlation value for the 448^th^ scanline was 0.376±0.046. Next, we applied 2 depolarizing stimuli at 200μA at a rate of 2 Hz and identified the most intensely responding 30 μm region (“hotspot”) within the field for both jRGECO1a and jGCaMP8f. On average, jRGECO1a and jGCaMP8f “hotspots” were separated by a distance of 182.2±21.1 μm, and never less than 107.5 μm (Figure 2E-2Ei). From this, we conclude that the robust spectral separation exists between the 535nm and 590nm channels, and that fluorescent structures in each channel are likely to represent transcriptionally distinct neuronal populations.

**Figure 2.**
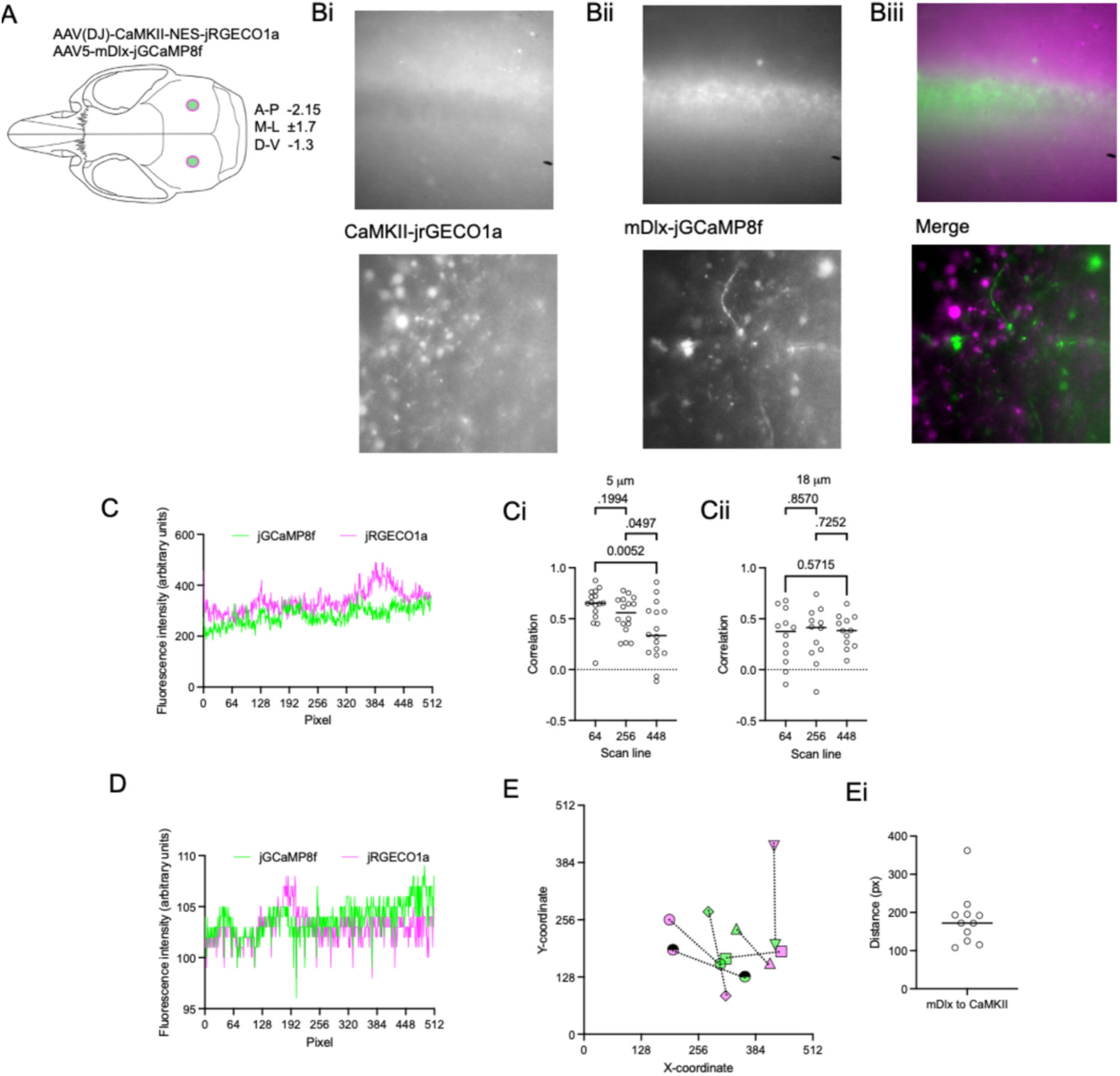
Tandem Ca2+ imaging of excitatory and inhibitory presynaptic terminals with robust spectral separation. A. Diagram of stereotactic coordinates for the injection of AAVs expressing jRGECO1a and jGCaMP8f under the control of the CaMKII promoter and mDlx enhancer for expression in excitatory and inhibitory neurons, respectively. Bi. Hippocampal CA1 regions expressing CaMKII-jRGECO1a fluorescence imaged in acute hippocampal slices at 40x in epifluorescence (upper panel) and in LLSM (lower panel). Bii. mDlx-jGCaMP8f fluorescence imaged in the same field at 40x in epifluorescence (upper panel) and in LLSM (lower panel). Biii. Merged dual-channel image with CaMKII-jRGECO1a in magenta and mDlx-jGCaMP8f in green. C. Representative trace of 595nm jRGECO1a fluorescence(magenta) and 535nm jGCaMP8f fluorescence(green) as a function of each pixel in a given scanline in the x-direction, at a slow acquisition rate of 2Hz. Ci. Scatterplot of orrelation of 595nm and 535nm fluorescence intensity for the 64^th^, 256^th^, and 448^th^ scanline in the image at an imaging depth of 5 μm into the slice. Cii. Scatterplot of correlation of 595nm and 535nm fluorescence intensity for the 64^th^, 256^th^, and 448^th^ scanline in the image at an imaging depth of 18 μm into the slice. D. Representative trace of 595nm jRGECO1a fluorescence(magenta) and 535nm jGCaMP8f fluorescence(green) as a function of each pixel in a given scanline in the x-direction, at the typical acqusition rate of 94Hz. E. Map of jRGECO1a (magenta) and jGCaMP8f hotspot Cartesian coordinates within six fields. Ei. Scatterplots of distance between the most intense jRGECO1a and jGCaMP8f hotspots in each field. The dataset represents 12-16 fields from 8 animals. Error bars represent means + S.E.M. P-values tabulated from unpaired Student’s t-test.

### Quantification of electrically evoked mDlx-jGCaMP8f and CaMKII-jRGECO1a fluxes

Because many hotspots can be identified in each of the two channels within a given 51×51 μm field, and many fields can be imaged in one experiment, we built a robust data analysis suite to automate and massively parallelize the process of finding hotspots and correcting for photobleaching/photoswitching artifacts. 600-image stacks of 535nm or 595nm fluorescent structures (Figure 3A-3B) acquired at a rate of 94 Hz were analyzed via automated ROI analysis (Figure 3Ai-3Bi, see Protocols 7: Data Analysis). During acquisition, two depolarizing electrical stimuli were administered a rate of 2 Hz. An internal p correction algorithm was used to remove photobleaching artifacts. Heatmaps of the most intensely responding puncta within each channel were created from the automated ROI analysis (Figure 3Aii-Bii). Fluorescence intensity changes subsequent to stimuli are plotted in Figure 3C and Figure 3D. From this, we have determined that the BLIMPS methodology is able to identify responding regions along GECI-labeled axons, and that the software suite is adequate to automate the analysis of large numbers of imaged fields.

**Figure 3.**
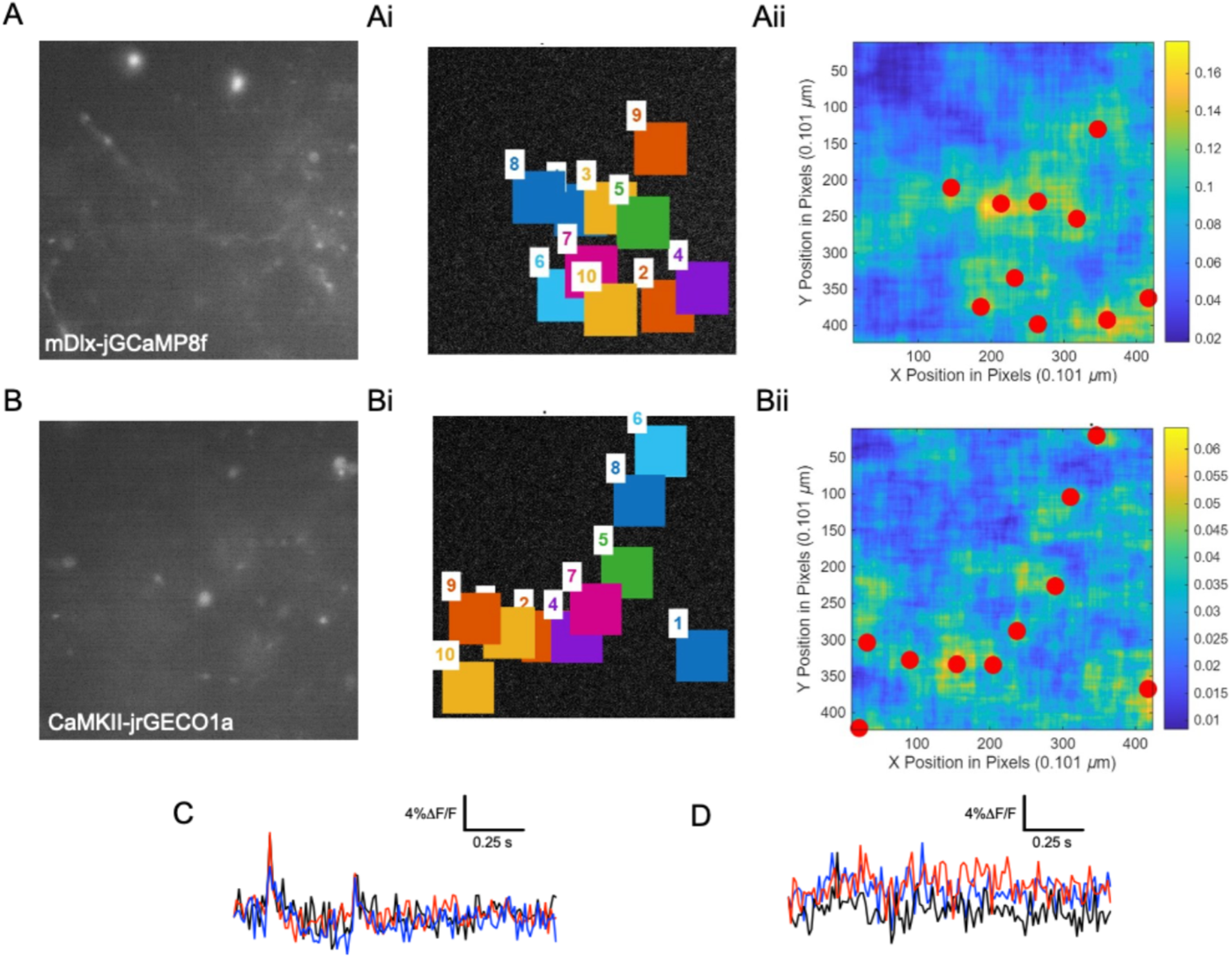
Automated analysis of electrically evoked tandem Ca^2+^ fluxes. A. 51×51 μm image of mDlx-jGCAMP8f+ axonal structures in CA1 dendritic fields imaged using BLIMPS as a 600-image time series at an acquisition rate of 94 Hz. Ai. Candidate regions of interest highly responsive to electrical stimuli in (A), as identified by spot intensity algorithms in the BLIMPS software suite. Aii. Heatmap of highly responsive regions in (A). B. 51×51 μm image of CaMKII-jRGECO1a+ axonal structures in CA1 dendritic fields imaged using BLIMPS as a 600-image time series at an acquisition rate of 94 Hz. Bi. Candidate regions of interest highly responsive to electrical stimuli in (B), as identified by spot intensity algorithms in the BLIMPS software suite. Bii. Heatmap of highly responsive regions in (B). C. 3 representative mDlx-jGCaMP8f fluorescence traces plotted in red, blue, and black, identified in (Aii). D. 3 representative CaMKII-jRGECO1a fluorescence traces plotted in red, blue, and black, identified in (Aii).

## MATERIALS

### AAV Injection

Mouse should be anaesthetized with isoflurane (Covetrus 11695067772) delivered viavaporizer calibrated to 2.7%. AAV delivered using 10uL Gas-tight injection system (WPI NANOFIL) with beveled 35G needle (WPI NF35BV), filled with AAV-CamKIIa-NES-jRGECO1a (SignaGen SL116372)and mDlx-jGCaMP8f (Vectorbuilder).

### Cutting Solution

93mM N-Methyl-D-glucamine (Sigma-Aldrich 66930), 2.5mM KCl (Sigma-Aldrich P9541), 1.2mM NaH2PO4 (Sigma-Aldrich S8282), 20mM HEPES (Fisher Scientific BP310), 10mM MgSO4 (M1880), 0.5mM CaCl2 (Sigma-Aldrich C4901), 25mM D-glucose (Fisher Scientific D16-500), 5mM ascorbate (Sigma-Aldrich 11140), and 3mM pyruvate (Sigma-Aldrich P8574). Solution should beoxygenated with 95% O2 and 5% CO2, with its pH adjusted to 7.4.

Frenguelli’s Modification of Ringer’s ACSF (FMRA): 124mM NaCl (Fisher Scientific S271), 2mM NaHCO3 (Fisher Scientific S233), 1.25 mM NaH2PO4 (Sigma-Aldrich S8282), 3mM KCl (Sigma-Aldrich P9541), 9mM D-glucose (Fisher Scientific D16-500), 1mM D-ribose (Thermo Scientific AC13236), 5μM adenine (Thermo Scientific AAA14906), 2 mM CaCl2 (Sigma-Aldrich C4901), and 1 mM MgCl2 (Sigma-Aldrich M2670). Solution should be oxygenated with 95% O2 and 5% CO2.

### Slice Preparation

Hippocampus slices are produced using a LeicaVT1200S vibratome (Leica Biosystems) and stainless steel blades (Campden Instruments 752-1-SS). When mounting slices to the LLSM, slices should be secured to glass coverslips (Fisherbrand 12-541) cut to size with Vetbond (3M 1469SB). Glass coverslips should first be affixed to the PMMA chamber with vacuum grease (Dow Corning 31735228).

### Pharmacological reagents

AMPAR antagonist CNQX (HelloBio HB0205) stored as10mM aliquots at −20°C, used at a final concentration of 5μM in FMRA. NMDAR antagonist AP5 (HelloBio HB0252) stored as 50mM aliquots at −20°C, used at a final concentration of 50μM in FMRA

## PROTOCOL

### 1. Startup

Power on and turn the keys of 488nm, 560nm, 642nm, and 532nm MPBC Continuous Wave Fiber Lasers. Turn on the A.M.P.I Master-8 stimulator, Scientifica electrode manipulator, table air compressor, Acopian linear power supply, X- and Z-galvo controllers, AA Opto-electronic D66 multipurpose digital synthesizer, Forth Dimension Displays SXGA-3DM monochrome SLM, Stanford Research Systems SIM900 Mainframe, and both Hamamatsu ORCA-Flash 4.0 V3 cameras.

With the stage out from under the lenses, reference the stage using the reference switch on PIMikroMove. This should bring the stage to a maximal Z position of 18.5000mm. Once finished, lower the stage down to 3.5000mm so it can fit under the lenses without colliding.

Open one GUI instance for each fiber laser and turn on the 488nm, 642nm, and 560nm lasers. Open and run the LLS software on LabVIEW.

### 2. Alignment of lattice light sheet

#### Alignment of lattice light sheet

The lattice sheet is aligned using standard procedures following placement of a fluorescein dye droplet between the excitation and emission lenses of the LLSM over a a clean PMMA chamber on the stage and move it in so the excitation and emission lenses are centered over chamber. After alignment the stage is withdrawn to allow sample placement. Bring the stage down to 3.5000mm. Remove the fluorescein dye from the chamber and rinse the lenses with ultrapure water.

### 3. Acute hippocampal slice preparation

Anaesthetize the mouse with isoflurane. Identify the xyphoid process via palpation and make a lateral incision below it to access the abdominal cavity. Slice through the diaphragm and ribcage to expose the heart. Insert the needle into the left ventricle then slice the right atrium. Perfuse the animal with approximately 20mL of ice-cold cutting solution until the liver becomes pale.

Decapitate the animal. Remove the skin from the skull. Insert the scissors into the central canal of C1 and cut in a rostral direction to the zygomatic arch on each side. Peel back the skull and transfer the brain into a dish of ice-cold cutting solution (see materials). Using a scalpel, separate the brain into two hemispheres along the sagittal axis. For improved friction, place each hemisphere on squares of filter paper. Remove the cerebellum and thalamus from the cortical structures.

Isolate intact hippocampi from adjacent parenchyma using a scooping motion with a curved metal spatula. Excise approximately 1mm of tissue from each end of the septotemporal axis. Mount the remainder of the hippocampus by affixing the temporal axis to the specimen disk with commercial cyanoacrylate adhesive. Attach the specimen disk to the magnetitic holder within a buffer dam filled with oxygenated ice-cold cutting solution. Secure a razor blade to the blade holder using an Allen wrench. Use blades no more than twice to minimize tissue damage.

Set the desired beginning and end positions for the slicer anterior and posterior to the tissue in the X-direction. Position the blade approximately 1 mm above the tissue in the Z-direction. Use the vibratome to slice individual 300 μm sections at a rate of 0.20mm/s, and 1.2mm amplitude. Immediately transfer slices into oxygenated ACSF containing the RibAde modification proposed by group of Bruno Frenguelli^26^ (termed Frenguelli’s modification of Ringer’s ACSF) (see materials).

Acute slices may be incubated at 34 C for 30 minutes post-slicing to facilitate the removal of dead cells. Hippocampal slices remain responsive and viable for up to six hours post-sacrifice.

### 4 Preliminary characterization of acute slices

LLS *ex vivo* biosensor imaging is contingent on fluorophore brightness: dim/poorly expressed fluorophores may be difficult to image. Prior to imaging acute slices, it is advised to verify biosensor expression and tissue viability using traditional fluorescence microscopy. Multiple types of instruments, including widefield/epifluorescence microscopes, confocal microscopes, and multiphoton systems may all be suitable for preliminary characterization: here, the simplicity and speed of the widefield system make it preferable for biosensor imaging.

Transfer the slice to the widefield imaging chamber gently using a Pasteur pipet. Secure the slice in position with a weighted harp, taking care to avoid damage. Check for the presence of jGCaMP8f+ and jrGECO1a+ fluorescent structures under blue and green LED illumination throughout the tissue, with appropriate filters. Position a twisted-pair stimulating electrode into contact with the surface of the slice upon the labeled axons within the region of interest. Bring relevant synaptic structures into focus at 40x magnification.

Program the Master-8 pulse stimulator and connected stimulus isolator to deliver 1-5 stimuli at frequencies appropriate for the experiment. To induce action potentials in CA1 dendritic fields, it is recommended to use a duration of 50-200 μs and stimulus amplitudes of 25-400 μA. Stimulus amplitudes and durations may vary depending on electrode material and the characteristics of the tissue.

Using the multi-dimensional acquire feature of MicroManager, capture a time series of electrically evoked 535nm jGCaMP8f images and 595nm jRGECO1a fluorescent images while applying electrical stimuli. After imaging, use the Plot Z-Axis profile function of ImageJ to assess DF/F changes contemporaneous with stimulus application.

To differentiate between electrically evoked Ca^2+^ fluxes and movement artifacts induced by electrostatic forces, it is advised to attempt to pharmacologically block evoked transients with well-characterized inhibitors, such as Cd^2+^. Manipulation of the Ca2+ concentration within the FMRA should alter the evoked transient size: smaller transients should be seen at 0.5mM Ca^2+^ relative to 2mM Ca^2+^, while movement artifacts will be invariant to external Ca^2+^ changes.

### 5. Mounting slices for use in the LLS system

Affix a 5 by 22 mm glass coverslip to the PMMA chamber using vacuum grease. Remove excess grease with a delicate task wiper. Transfer the slice to the chamber gently with a Pasteur pipet under oxygenated FMRA. Position the slice on the coverslip so an electrode can be placed without covering the region of interest.

Secure the slice to the coverslip using 0.25uL-drops of butyl cyanoacrylate at the edges of the tissue and let the drops dry before moving the dish. Verify that the slice is secure by gently shaking the dish. Avoid depositing butyl cyanoacrylate on or around the anatomical region of interest to prevent tissue warping.

With the stage lowered, transfer the chamber to the rim of the stage. Affix the inlet and outlet of the fluidics to the chamber with paraffin wax. Start the flow of oxygenated FMRA immediately.

Check the temperature of the bath using the thermocouple and verify it is in the physiologically appropriate range for your sample. Position the chamber so the slice will fall between the excitation and emission objectives.

Using the dissecting microscope and the Scientifica electrode manipulator, place a twisted-pair stimulating electrode on the surface of the slice and make ensure that contact between the electrode and the lens is avoided. Position the stage so that the slice is centered beneath the excitation and emission objectives. Turn on the illumination and slowly raise the stage until both lenses are just immersed in FMRA. Once immersed, raise the stage at 0.2mm intervals to look for the surface of the tissue. When the surface of the slice is in focus, soma and neuropil should be visible.

### 6. LLSM imaging of single presynaptic terminals

Illuminate with the 560nm laser and look for the bright yellow-green laser dot at or near the surface of the tissue. Bring the stage to the desired position in the x and y directions. Observe the position of the laser dot relative to the electrode tip to find fields of view in proximity to the electric field induced by the point charge from the electrode.

The electric field may not induce action potentials within the field of view if the electrode tip is embedded too deep in the tissue. If so, raise the electrode to the surface of the tissue. If the electrode is in contact with the tissue, moving the electrode arm gently should move the tissue.

With the terminus of the electrode at the edge of the field of view at 2048×2048px and the surface of the neuropil in focus, turn off the illuminator and activate the 488nm laser. Structures fluorescent at 535nm should be faintly visible with an exposure time of 10ms. Cut the field of view to 512×512px. If the lattice is positioned at the surface of the tissue, blebbed structures of dead tissue from the slicing process should be visible on the right side of the field. Move the stage closer to the objectives slowly in 0.1-0.5-micron increments until live soma and axons are in view.

Program the Master-8 pulse stimulator and connected stimulus isolator to deliver 1-5 stimuli at frequencies appropriate for the experiment. A delay of at least 2 s is recommended to mitigate initial jGCaMP8f and jRGECO1a photoswitching.

Select an appropriate filter from the emission filter wheel, either the Chroma 520/40m for jGCaMP8f or the Semrock Brightline 593/40 for jRGECO1a. For high-speed imaging, sequential imaging of jGCaMP8f and jRGECO1a is recommended, as the filter wheel is unable to cycle between the two.

With the pulse stimulator connected to the camera using TTL logic, begin image acquisition using the Z-stack function. Set the Z-galvo and Z-piezo range to 0 for imaging a single section 0.5 μm in height, with 0 seconds between stacks. Each frame in the time series is treated as a singular stack consisting of one image.

Save the Z-stack using the disk icon to the right of the camera display window.

After identification of responsive fluorescent puncta in ImageJ, VGCC currents localized to axons can be pharmacologically isolated via the application of CNQX to block Ca^2+^-permeable AMPA and kainate receptors along with AP5 to block NMDA receptor currents.

At the conclusion of imaging, pass ultrapure water through the fluidics of the instrument for 20 minutes while the lenses are still immersed solution followed by air for 10. Rinse the PMMA chamber out with ultrapure water and discard the slice and coverslip. Carefully dry the excitation and emission objectives using lens paper with a “flossing” motion. The instrumentation can be powered down between uses.

### 7. Data Analysis

The BLIMPS technique includes an open-source data analysis package available on the Alford Lab GitHub at github.com/AlfordLab/BLIMPS.

Fluorescence traces were processed in MATLAB and ImageJ, where regions of interest were manually defined and time-dependentvalues were extracted. These normalized signals provided a quantitative measure of presynaptic calcium dynamics.

Raw lattice light-sheet datasets underwent a structured post-processing pipeline designed to preserve the fast kinetics of jGCaMP and jRGECO signals while minimizing motion-related artifacts. Image sequences were first corrected for lateral drift using subpixel registration algorithms optimized for high-frame-rate data. Following stabilization, regions of interest (ROIs) were manually or semi-automatically segmented to isolate defined structures, including dendrites and axonal boutons. Fluorescence time-series were extracted from each ROI to generate raw intensity traces. Automated ROI detection was performed using a structured image-processing pipeline designed to isolate biologically meaningful calcium-active structures while minimizing noise and segmentation artifacts. The workflow began with watershed-based segmentation, in which *imregionalmax* was applied to identify local fluorescence maxima within the calcium imaging stack. These maxima typically correspond to the centers of active cells or subcellular compartments. The resulting binary mask was then labeled using *bwlabel* to generate discrete candidate ROIs. To refine these candidates, an intensity-based threshold was applied to the maximum-projection image. This thresholding step restricted ROI detection to regions exhibiting significant calcium-dependent fluorescence changes, thereby excluding background fluctuations and low-signal structures unlikely to represent true biological activity. Following thresholding, ROIs were filtered by size to remove spurious detections. Very small regions, often arising from photon noise or single-pixel fluctuations, were excluded, as were abnormally large regions that typically reflect motion artifacts, merged cells, or out-of-focus fluorescence. This size-based filtering step ensured that the final ROI set consisted of well-defined cellular or subcellular structures suitable for quantitative calcium-signal extraction. Together, these automated segmentation procedures produced a high-quality ROI map optimized for downstream kinetic and spatial analyses of jGCaMP and jRGECO signals.

Each trace was normalized to its baseline fluorescence F_0_, calculated from a pre-stimulus interval, to obtain ΔF/Fvalues suitable for quantitative comparison across indicators and preparations. Adouble exponential decay baseline was subtracted from the ΔF/Fvalues to flatten the signal and to simplify the peak fitting routine which involves summing these shapes with a baseline signal. The custom multi-peak stimuli fitting function combines a Gaussian rise and double exponential decay. To create a peak with a Gaussian rise (left side) and a double exponential decay (right side/tail), the following model is often used, where the peak is split at its centroid (*x*_0_):

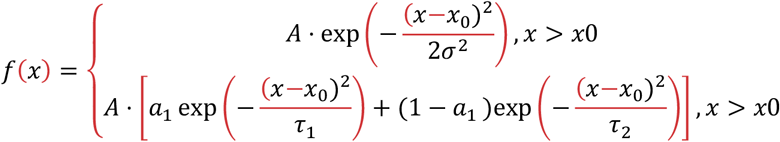

Parameters

*A* : Amplitude (peak height)

- *x*0 : Center of the peak
- σ : Gaussian width (controls rise speed)
- τ_1_, τ_2_: Decay time constants (fast/slow decay)
- *a*_1_: Weight of the first decay component

Because this function is nonlinear and requires finding optimal parameters for multiple peaks, we used lsqcurvefit in MATLAB.

From these normalized and baseline corrected traces, several kinetic parameters were computed. Peak amplitude captured the maximal calcium-dependent fluorescence change. Rise time (τ_rise_) was defined as the interval required for the signal to increase from 10% to 90% of its peak, providing a measure of activation kinetics. Decay time (τ_decay_) quantified the return from peak to 50% of peak amplitude, reflecting calcium clearance dynamics. The area under the curve (AUC) was integrated over the event duration to estimate total calcium load.

## DISCUSSION

Here, we have developed a novel methodology to assess presynaptic activity in two transcriptionally distinct cell populations in an *ex vivo* tissue preparation. The BLIMPS technique offers numerous functional advantages relative to current techniques. First, with axial resolution of only 0.81 μm, BLIMPS offers unparalleled optical sectioning capability to trace activity to single anatomical features within a preparation at high speeds without deconvolution. Our implementation also offers robust lateral resolution of 0.501 μm, sufficient to identify structures such as individual synaptic terminals. Together, this represents much better spatial resolution compared to widely adopted multiphoton methodologies^27^, with imaging rates of 94 Hz routinely achieved. The bleaching of fluorophores and photodamage to living tissue is minimal relative to comparable multiphoton techniques, which can heat tissue 1-2° C at or below the illuminated region throughout the time course of an experiment^28^. Critically, the technique permits the implementation of multiple biosensors with distinct excitation/emission spectra using rapid sequential imaging. The technique should have forward compatibility with newly developed genetically encoded biosensors with unique excitation/emission spectra such as FR-GECO1^29^, requiring only unique emissions filters. The use of AAVs to deliver genetically encoded biosensors enables a wide range of genetic disease models and environmental stressors such as maternal immune activation^30,31^ to be used with BLIMPS. One potential downside of BLIMPS is the considerable cost and technical complexity of building the LLSM.

The implementation of red fluorescent biosensors with ∽590nm fluorescence subsequent to excitation with the 560nm laser line presents technological challenges to accompany experimental possibilities. The reduced sensitivity and slower kinetics of jRGECO1a make it less useful for assessing neuronal calcium buffering relative to the jGCaMP8 series of fluorophores: it is anticipated that the next generation of red indicators such as the mScarlet-derived SCaMP will bridge this functional gap. Upon availability, these indicators can be substituted for jRGECO1a with minimal friction. Outside of Ca^2+^ sensing, recently developed red or far-red biosensors for dopamine^32,33^ and serotonin^34^ can readily be incorporated into our workflow with jGCaMP8, allowing simultaneous assessment of dopaminergic and serotonergic neurotransmission in tandem with either excitatory or inhibitory neuronal activity. Conversely, fast and responsive green fluorescent biosensors like iGluSnFr4^35,36^ or GRAB-NE can be imaged here in tandem with jRGECO1a, an approach that was previously utilized in other imaging and photometry modalities such as dual-NeCa^37^.

Imbalances between excitation and inhibition within distinct neuronal circuits is thought to contribute to the pathology of number of diseases, such as Alzheimer’s disease^1,2^ and epilepsy^38^. Excitatory neurotransmission is accomplished via glutamatergic neurons, whereas inhibitory neurotransmission occurs through GABAergic neurons. These cell types are transcriptionally distinct and emerge from different developmental lineages^39,40^, enabling the selective labeling of each with cell-type specific promoters directing ORF transcription within an AAV. The use of genetically encoded biosensors with differential fluorescence in response to changes in cytosolic Ca^2+^ is generally accepted as a proxy for neuronal activity states^24,41^: the measurement of Ca2+ fluxes in two populations of neurons within a given preparation can represent activity ratios such as E-I balance. Because of its fast kinetics and robust spatial resolution, the BLIMPS technique is one of the only methodologies suitable to assess the E-I balance at a given population of presynaptic terminals within a user-defined circuit of interest and disease model. We utilized the well-studied 1.3kb fragment of the calcium/calmodulin-dependent protein kinase II promoter^42^ in tandem with the mDlx enhancer^43^ to drive expression of two spectrally separated genetically encoded calcium indicators specifically in excitatory pyramidal cells and inhibitory neurons within the hippocampus. As expected, minimal overlapping expression exists between CaMKII-positive labeled cells and mDlx-positive labeled cells (Figure 2). A positive correlation of approximately .35 was observed between the 595nm jrGECO1A and 535nm jGCaMP8f channels. While it may be anticipated that no correlation (a value of 0) would be observed with complete spectral separation, voids where no fluorescent material is present in either channel will produce the same amount of background signal in both channels, creating correlation. CaMKII-positive projections were observed extending into the stratum oriens and stratum radiatum, while mDlx-positive projections formed a mesh-like structure roughly ensheathing the stratum pyramidale (Figure 2Ai). No dual-positive soma were observed at any point, demonstrating the validity of our promoter selection, and highly responsive spots within fields were separated by considerable physical distances of hundreds of μm (Figure 2D), indicating that responsive jRGECO1a+ and jGCaMP8f+ projections originated from separate populations of excitatory and inhibitory neurons, respectively.

In conclusion, we have developed BLIMPS, a sensitive, kinetically robust, and high-resolution system for the imaging of presynaptic Ca^2+^ within multiple populations of neurons within a given *ex vivo* preparation. Because of its superior axial resolution and fast acquisition speeds ideal for the assessment of biosensor kinetics in multiple populations, BLIMPS represents a methodological advance for the study of 4D cellular physiology of presynaptic terminals, and may be implemented in a variety of neuronal or neuron-effector circuits. The BLIMPS software suite enables rapid analysis of large data sets, empowering experimenters with new capability to assess multidimensional population-level differences in synaptic activity.

## ACKNOWLEDGEMENTS

AAV-mDlx-jGCaMP8f-WPRE was cloned and prepared by VectorBuilder, Inc. Funding was provided by NIH K25AG086674 (Potcoava), Funding was provided by NIH 1K25AG086674 (Potcoava), R01NS111749 (Alford) and Canadian Institute for Health Research CIHR 507738 (Alford) We express gratitude to the research support/operations team at the Department of Anatomy and Cell Biology University of Illinois Chicago, including Ashley Miller, Aggeliki Gikas, Lea Smucker, and Kathy Hernandez-Chavez.

